# High-resolution single-cell sequencing of malaria parasites

**DOI:** 10.1101/130864

**Authors:** Simon G. Trevino, Standwell C. Nkhoma, Shalini Nair, Benjamin J. Daniel, Karla Moncada, Stanley Khoswe, Rachel L. Banda, François Nosten, Ian H. Cheeseman

## Abstract

Single-cell genomics is a powerful tool for determining the genetic structure of complex communities of unicellular organisms. Patients infected with the malaria-causing parasite, *Plasmodium falciparum*, often carry multiple, genetically distinct parasites. Little is known about the diversity and relatedness of these lineages. We have developed an improved single-cell genomics protocol to reconstruct individual haplotypes from infections, a necessary step in uncovering parasite ecology within the host. This approach captures singly-infected red blood cells (iRBCs) by fluorescence-activated cell sorting (FACS) prior to whole genome amplification (WGA) and whole genome sequencing (WGS). Here, we demonstrate that parasites in late cell cycle stages, which contain increased DNA content, are far superior templates for generating high quality genomic data. Targeting of these cells routinely generates near-complete capture of the 23Mb *P. falciparum* genome (mean breadth of coverage 90.7%) at high efficiency. We used this approach to analyze the genomes of 48 individual cells from a polyclonal malaria infection sampled in Chikhwawa, Malawi. This comprehensive dataset enabled high-resolution estimation of the clonality and the relatedness of parasite haplotypes within the infection, long-standing problems in malaria biology.

## Background

In recent years, single-cell genomics has helped unravel the population dynamics of unicellular organisms [1-4], cancer cells [5] and developmental lineages [6] in multicellular organisms. Efforts to eradicate malaria, the world’s most consequential parasitic disease, can greatly benefit from understanding the genetic strategies that enable *Plasmodium falciparum* communities to persist and thwart interventions. For instance, genome sequencing has played a pivotal role in understanding the spread of drug-resistant parasites [7], global structuring of parasite populations [8] and selection of vaccine candidate loci [9]. In locations where malaria is endemic, patients are often infected with multiple parasite lineages [10-14]. Basic details about the genetic structure of individual malaria infections, such as the number of distinct parasite lineages, their diversity and their relationship to one another, remain unclear in polyclonal infections. One promising strategy for dissecting such complicated population structures is to construct complete, individual haplotypes as has been performed to understand the evolution of cancer cells [5].

Current methods to understand the complexity and diversity of malaria infections are generally reliant on using DNA extracted from patient blood to make indirect inferences from either PCR genotyping or whole genome sequencing (WGS). Using such bulk DNA approaches, the number of haplotypes within an infection can be inferred either through maximum likelihood estimation from PCR genotyping [8, 15-17] or through unfixed reads in WGS data [8, 18-20], and progress has been made in computationally deconvoluting haplotypes from WGS data using methods analogous to phasing diploid human sequences [21]. While these approaches are scalable, affordable, and can provide estimates of key demographic parameters they are generally reliant on assumptions such as random mating [15], which are frequently violated in parasite populations. Thus, they methods are unable to provide accurate haplotype-level resolution of infections. Consequently, direct methods to understanding the complexity of infections can add considerable depth and accuracy as has been the case for direct phasing methods of human genomes [22].

Two methods have been developed to experimentally isolate individual haplotypes from polyclonal infections: culture-based dilution cloning [23] and single-cell genomics [3]. Dilution cloning can generate high-quality bulk DNA from clonal expansion of individual parasites [12] but is labor intensive, prone to contamination and is reliant on parasites to thrive in culture, making it inappropriate for large-scale experiments. As a response to these limitations, we previously developed a single-cell genomics approach based on fluorescence-activated flow cytometry (FACS) and whole genome amplification (WGA) [3]. In this design, infected red blood cells (iRBCs) were cultured overnight and subsequently stained with a fluorescent DNA dye, which allows isolation of individual iRBCs by FACS. WGA of lysed single-cells enabled sufficient DNA for WGS or targeted genotyping applications. Unfortunately, a high rate of allelic and genomic dropout limited the analyzable proportion of the genome to less than 50% and required costly quality control steps to be done prior to library preparation and WGS.

The efficiency of successful WGA and breadth of genomic coverage are major technical barriers for the broad-scale analysis of samples. Recent work towards improving WGA of human nuclei [24] and studies of bulk *P. falciparum* WGA [25] suggest that initiating amplification with more than a single genome copy would generate better quality data. During the ∼48 hour life cycle of *P. falciparum* in the blood, late-stage iRBCs generate an average of 16 clonal copies of the parasite’s genome by DNA replication [26]. We hypothesized that these DNA-rich parasites would provide multiple templates for WGA, thereby improving the reaction and subsequent WGS data quality.

We found that in an asynchronous culture of a well-characterized *P. falciparum* laboratory line, HB3, cells with the highest DNA content yield near-complete genome coverage (mean 92.4%). After optimizing our protocol in clinical samples, we interrogated a polyclonal infection (MAW0) from Chickhwawa, Malawi and recovered similarly high genome coverage (mean 90.7%) for 48 out of 48 attempted reactions. These data allow fine scale estimation of diversity and relatedness within a single malaria infection.

## Results and Discussion

### iRBCs that contain high DNA content are superior targets for WGS

In the 48 hour blood stage of the malaria lifecycle, parasites are haploid and in the earliest stage of their life cycle contain only a single copy of the genome. By the latest life cycle stage, they harbor ∼16 copies prior to bursting and invasion of new RBCs. To test whether targeting late-stage iRBCs would generate gains in genome data quality, we performed WGA of parasites with increasing DNA content. A commonly used laboratory-adapted line, HB3 (MR4, VA), was thawed and cultured for several weeks to allow asynchrony in cell cycle progression. Parasite DNA in the cultured cells was stained with Vybrant DyeCycle Green (Fig. 1a) and analyzed by fluorescence-activated flow cytometry. Three fluorescent subpopulations of DNA-containing cells were observed by flow cytometry, reflecting the asynchronicity of the culture. Event gates capturing these three populations were drawn, denoted low (L), medium (M) or high (H), based on increasing levels of fluorescence due to increasing DNA content (Fig. 1b).

**Fig. 1.**
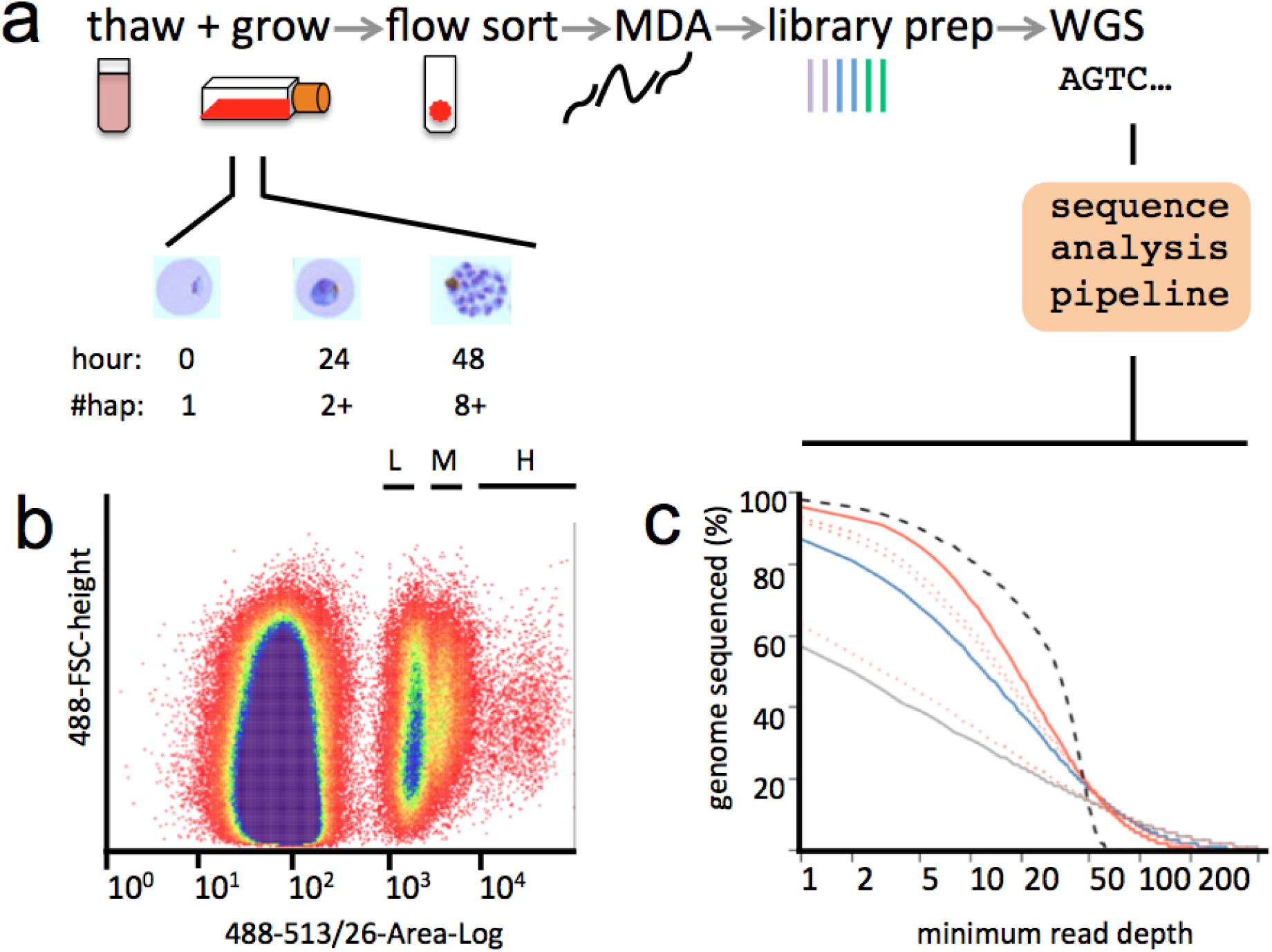
Targeted single-cell genomics of late-stage malaria parasites. (a) Cryopreserved iRBCs are thawed and grown under standard conditions for 40 hours, generating late-stage parasites with multiple genome copies. DNA-stained iRBCs are sorted into individual tubes by FACS. To generate high quality reactions, high DNA content, late-stage parasites in the H gate are freeze-thaw lysed prior to MDA, library preparation and WGS. (b) An asynchronous culture of HB3 containing parasites with different amounts of DNA. The x-axis shows the fluorescence intensity, and the y-axis the size of each cell. (c) Genome coverage obtained by sequencing cells from the L, M and H gates. The plot shows the proportion of the genome (y-axis) sequenced to at least a given minimum read depth (x-axis). The black dashed line is data obtained by routine sequencing of high quality DNA from a laboratory derived line. The solid lines denote cells from the L (grey), M (blue) and H (red) gates, with dotted red lines additional cells from the H gate. All libraries were downsampled to 30X coverage for comparability.

Individual cells from each gate were sorted into single tubes, freeze-thawed, and WGA was carried out by multiple displacement amplification (MDA, REPLI-g Midi, Text S1). Stringent protocols were implemented to minimize the risk of contamination (Methods, Text S2). As an initial test for DNA quality, two parasite-specific genes, *pfcrt* and *dhfr* were amplified from HB3 WGA reaction products by standard end-point PCR, providing a qualitative assessment for genome amplification. Fifty percent (5/10) of reactions failed to yield product for cells in the L gate, compared to 20% (2/10) for cells in the M gate and 0% (0/14) for cells in the H gate (Fig. S1). We took this as a preliminary indication that dropout of alleles might occur less readily in MDA of iRBCs containing higher DNA content.

### Near-complete capture of malaria haplotypes

We next performed WGS of representative reactions from cells sorted in each gate. Two metrics were used to determine the usefulness of sequencing data: read purity (the fraction of observed reads that map to the *P. falciparum* reference) and genome coverage (the fraction of the genome with at least one read mapped). For the HB3 cells (three cells total, one cell from each gate), > 87% reads mapped to the *P. falciparum* reference genome in every gate, demonstrating that our guidelines for sterility were sufficient to eliminate outside contamination. Interestingly, the HB3 L gate reaction was marked by moderate genome coverage (64.8% coverage) while cells sorted by the M (93.6% coverage) and H gates (97.4% coverage) yielded high coverage, similar to the genome coverage recovered from bulk DNA (97.8%) (Fig. 1c, Table S1). Subsequent sequencing of three additional cells from the H gate confirmed high capture of the parasite genome in two out of three cells (95.6%, 95.2%, 71.5%).

**Table 1.**
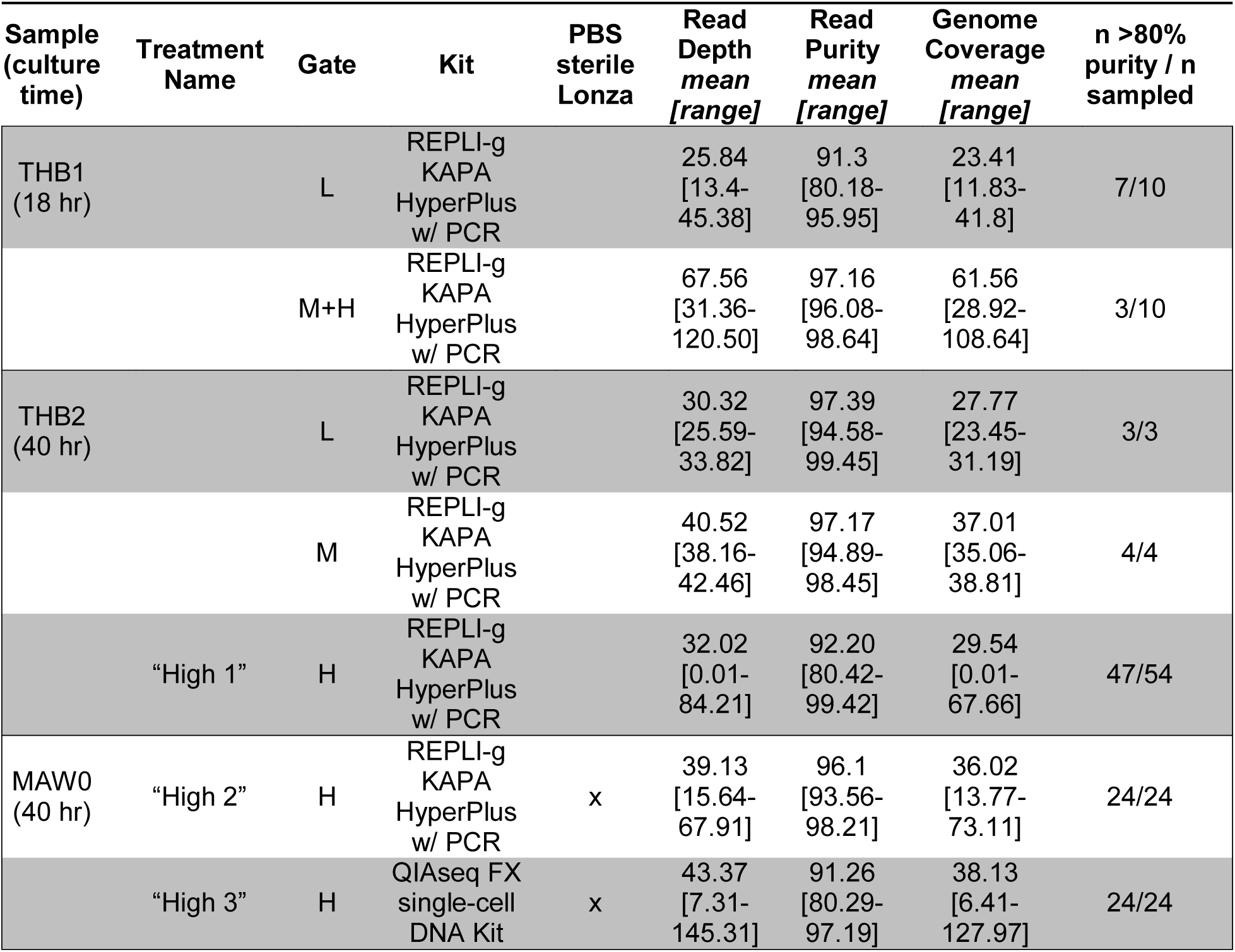
Summary of method parameters and data quality statistics for clinical samples. Only samples with read purity > 80% were included in this dataset.

| Sample<br>(culture<br>time) | Treatment<br>Name | Gate | Kit | PBS<br>sterile<br>Lonza | Read<br>Depth<br>mean<br>[range] | Read<br>Purity<br>mean<br>[range] | Genome<br>Coverage<br>mean<br>[range] | n >80%<br>purity / n<br>sampled |
| --- | --- | --- | --- | --- | --- | --- | --- | --- |
| THB1<br>(18 hr) |  | L | REPLI-g<br>KAPA<br>HyperPlus<br>w/ PCR |  | 25.84<br>[13.4-45.38] | 91.3<br>[80.18-95.95] | 23.41<br>[11.83-41.8] | 7/10 |
|  |  | M+H | REPLI-g<br>KAPA<br>HyperPlus<br>w/ PCR |  | 67.56<br>[31.36-120.50] | 97.16<br>[96.08-98.64] | 61.56<br>[28.92-108.64] | 3/10 |
| THB2<br>(40 hr) |  | L | REPLI-g<br>KAPA<br>HyperPlus<br>w/ PCR |  | 30.32<br>[25.59-33.82] | 97.39<br>[94.58-99.45] | 27.77<br>[23.45-31.19] | 3/3 |
|  |  | M | REPLI-g<br>KAPA<br>HyperPlus<br>w/ PCR |  | 40.52<br>[38.16-42.46] | 97.17<br>[94.89-98.45] | 37.01<br>[35.06-38.81] | 4/4 |
|  | "High 1" | H | REPLI-g<br>KAPA<br>HyperPlus<br>w/ PCR |  | 32.02<br>[0.01-84.21] | 92.20<br>[80.42-99.42] | 29.54<br>[0.01-67.66] | 47/54 |
| MAW0<br>(40 hr) | "High 2" | H | REPLI-g<br>KAPA<br>HyperPlus<br>w/ PCR | x | 39.13<br>[15.64-67.91] | 96.1<br>[93.56-98.21] | 36.02<br>[13.77-73.11] | 24/24 |
|  | "High 3" | H | QIAseq FX<br>single-cell<br>DNA Kit | x | 43.37<br>[7.31-145.31] | 91.26<br>[80.29-97.19] | 38.13<br>[6.41-127.97] | 24/24 |

Shortening the length of MDA reactions has been shown to improve the evenness of genome coverage by restricting runaway amplification in other contexts [28]. Thus, we sampled three reactions (L, M, and H gate) across several time-points (4.5, 8 and 16 hours) of WGA. Surprisingly, all single-cell reaction times yielded similar depth of coverage (Table S1, Text S1), suggesting amplification bias is minimal between 4.5 and 16 hours of reaction, perhaps due to diminishing enzyme activity or peculiarities of primer annealing in AT-rich genomes.

Previously, single-cell sequencing of malaria cell lines (HB3, 3D7) and clinical samples generated sequence data with variable genome coverage [3]. It is important to note that in thawed clinical blood samples, early-stage parasites are enriched because, unlike late-stage parasites, they are both present in circulation and able to survive cryopreservation. Our previous protocol used an overnight (18 hour) culture, after which parasites are unlikely to have progressed sufficiently far through the cell cycle to have undergone multiple rounds of DNA replication. We re-designed our protocol to enrich for late-stage parasites by analyzing two clinical samples collected on the Thai-Burmese border, grown either for 18 hours or 40 hours prior to FACS (THB1, THB2, respectively). Analysis of 25 single-cell DNA libraries from these samples showed that cells sorted from the H gate generated better genome data quality than L or M gate cells (Fig. S2, Table S1, Text S1), consistent with the trend observed for HB3 cells (Fig. 1). However, the breadth of genome coverage was lower than for laboratory-derived cells (L gate: mean=31.5% (range 22-41%), M gate: mean=50.0% (range 24-80%), H gate: mean=68.2% (range 54-96%)).

To further optimize our protocol for clinical samples, we made two further modifications prior to analyzing a third clinical sample collected in Chikhwawa, Malawi (MAW0). First, since amplification of small amounts of DNA is extremely sensitive to DNA contamination in reagents [29, 30], we elected to autoclave PBS (Lonza) used to “capture” sorted cells. Second, we processed half of the samples with a PCR-free library preparation method with the goal of lowering amplification bias introduced after MDA. In total, we processed 24 iRBCs using REPLI-g MDA and PCR-including KAPA HyperPlus library preparation and 24 iRBCs using the PCR-free QIAseq FX Single Cell DNA Library Kit (Qiagen). In this experiment, MAW0 was cultured *ex vivo* for 40 hours prior to sorting and only H gate iRBCs were analyzed.

All tested single-cell libraries from MAW0 were of high purity (mean reads mapped to the *P. falciparum* reference genome 93.7%, range 75.9-98.2%, Fig. S4). The proportion of the genome for which we were able to generate sequence data for was uniformly high across the 48 single-cells, with an average of 90.7% (range 52.4-98.6%) of the genome containing at least one correctly mapped read. This rose to an average of 92.3% (range 48.2-99.8%) of the genome after excluding highly polymorphic regions generally not amenable to most routine sequence analysis (Table S1). Furthermore, PCR-free library amplification improved the mean genome coverage and reduced sample-to-sample variation in genome coverage (Fig. 2, Table S1). To ensure that improvements in our protocol were not due to inadvertent capture of multiple cells, we analyzed the proportion of mixed base calls at high coverage (> 30X) sites. This resulted in the exclusion of 5/48 single cell sequences where > 5% of sites contained < 95% of reads supporting a single genotype. These were likely conservative thresholds as putatively clonal *P. falciparum* genome sequences can frequently contain unfixed base calls due to challenges in aligning to the highly AT-rich and repetitive reference genome [31].

**Fig. 2.**
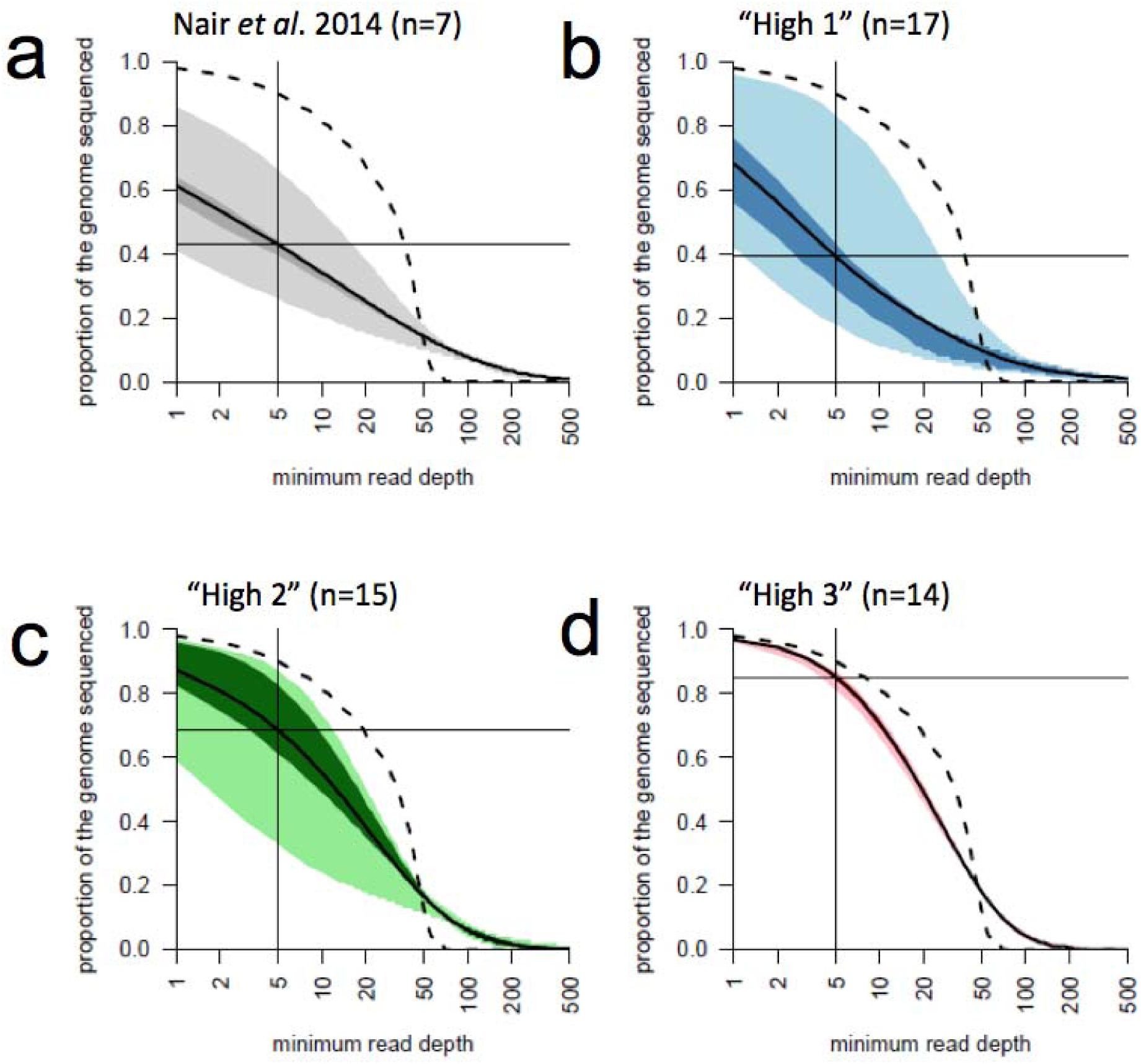
Comparison of genome coverage for single-cell WGS libraries. Each plot shows the same statistic as in Fig 1c, including the median value (solid line) with the interquartile range (dark shading), and the range (light shading). Genome coverage as a function of read depth from WGS data collected by previous work THB0 (a) or H gate-sorted cells grown for 40 hours from THB2 (b), MAW0 (c), (d). (a), (b) and (c) were processed by REPLI-g and KAPA HyperPlus library preparation with PCR amplification. (c) and (d) were sorted into sterilized PBS (Lonza) and (d) was processed with the QIAseq FX singlecell DNA Kit. All libraries were downsampled to 30X coverage for comparability.

Clear gains in single-cell data quality emerge when comparing the progress of our genome coverage through successive methodological improvements. We took the seven cells (THB0) sequenced using our previous protocol [3], and 46 single-cell sequences from the three treatments described here: (i) H gate with manufacturer’s PBS and KAPA HyperPlus with PCR Library Amplification Kit (“High 1”, THB2), (ii) same as *i* except for inclusion of autoclaved PBS sort “capture” buffer (Lonza) (“High 2”, MAW0) or (iii) same as ii except with QIAseq FX single-cell DNA Kit (“High 3”, MAW0). We randomly downsampled BAM files so each library had a mean of 30X coverage for comparability. Notably, cells processed by the original method [3] had been pre-selected as containing a high proportion of genotype calls from a larger panel of isolates, potentially overestimating the quality of this data. Fig. 2 shows the steady increase in data quality throughout the development of this method. We attribute these improvements to the targeting of late-stage parasites, using high quality, sterile reagents and omitting PCR amplification of library preparations. We have gathered similar single-cell genomic data quality using this approach on other clinical samples, suggesting MAW0 is not a unique case.

As is the case for human nuclei [24], we demonstrate that malaria parasites undergoing replication serve as better starting points for WGA. We hypothesize that the presence of multiple copies of the same alleles increases the chances that a given DNA segment will be successfully primed and amplified. However, other factors may also play a role in the accessibility of DNA to MDA reaction components, such as protein:DNA contacts and differences in membrane composition at different life cycle stages. These were concerns for malaria genomes, which are housed beneath several membranes and require both freeze-thaw and chemical lysis prior to MDA [3].

Additional gains in purity and target genome coverage in single-cell WGA might be attained by including malaria-specific primer sets [8], using exome capture, or optimizing UV treatment of reagents prior to WGA [32], though our observed read purity is sufficiently high for most downstream applications. Currently, Qiagen does not provide custom primer sets included in the REPLI-g kit, so care must be taken to prevent contamination if alternative primer sets are explored.

The MAW0 sample was collected from Chikhwawa, Malawi, an area of intense malaria transmission, where infected individuals are likely to contain many parasite lineages. This presented an excellent opportunity to test whether our optimized protocol could dissect the complexity of a potentially challenging, diverse infection (MAW0, described above). We examined two features of the data: 1) how well haplotypes from the infection are represented in single-cell genomic analysis and 2) the overall patterns of diversity and relatedness between parasites.

### Haplotypic Diversity

The number of unique haplotypes within an infection is a key measure of diversity. Estimating the number of haplotypes directly from single-cell data should be a simple task of counting the observed number of unique haplotypes (assuming the infection has been sampled comprehensively). A complication of this is that mutations in the parasite genome accrue during an infection at a rate of ∼1x10^-9^ per bp per replication cycle [33]. In addition, errors induced by sequencing WGA products can introduce differences between haplotypes. While most errors may be accounted for during genotype calling, this process is imperfect and true *de novo* mutations are often retained in high quality data sets.

Fig. 3a plots the number of unique haplotypes inferred against the proportion of pairwise SNP differences between single-cell sequences drawn from MAW0. The number of haplotypes estimated rapidly declines over low levels of pairwise differences, likely reflecting the exclusion of unique *de novo* mutations and sequencing and/or amplification-induced errors. With increasing pairwise difference, the estimation of haplotype number reaches a plateau at 7. Given a suitable estimate of mutations and error rates expected during single-cell sequencing (the error rate of our sequencing is ∼1x10^-7^ per base, the estimated error rate of WGA is 1.4x10^-5^ per base [34], 202 false positive mutations per 20Mb core genome sequence are expected (equating to 0.25% of the 19,713 SNPs called in MAW0 data). We suggest that a suitable threshold to collapse together individual sequences into shared haplotypes for MAW0 is 0.5% (0.25% differences per sequence) and shown by the vertical red dashed line in Fig. 3a. Beyond differences of > 10% between sequences the estimates rapidly collapse as genuine distinguishing variation is eliminated. This estimate of 7 distinct haplotypes is similar to previous estimates from single locus deep sequencing performed in Malawi [17].

**Fig. 3.**
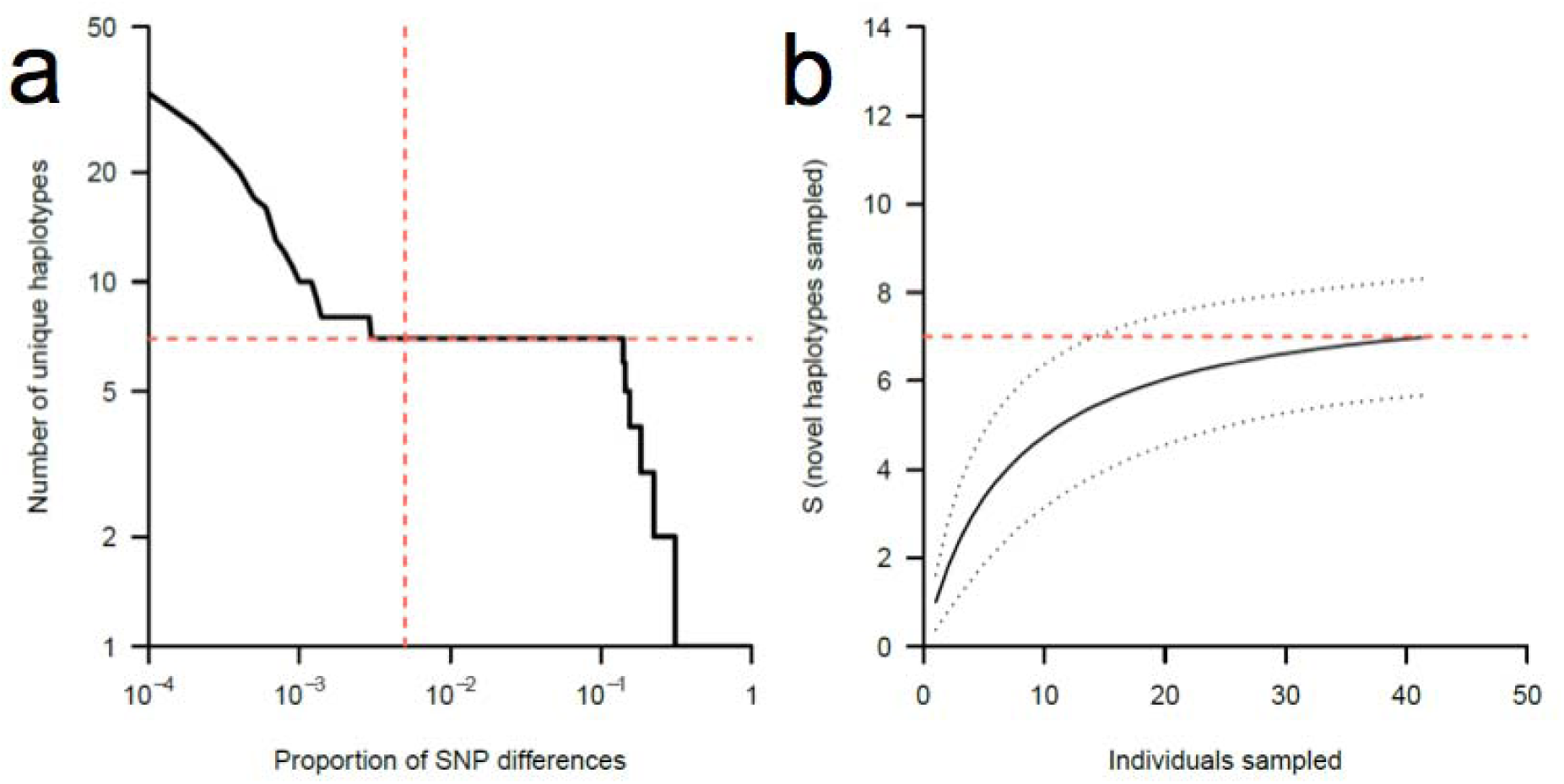
Estimation of the number of unique haplotypes in a complex infection. The number of unique haplotypes observed in MAW0 using an increasingly permissive threshold for pairwise differences (a). The vertical red line shows the point at which we estimate few errors will define new haplotypes while the horizontal red line shows the estimated number of haplotypes at this threshold. Rarefaction curve for 43 single cells from the MAW0 infection, 95% confidence interval in dashed black line (b). The red dashed line is the estimated number of haplotypes from (a).

In order to determine whether or not the infection had been sampled to an appropriate depth, rarefaction analysis [35] was performed on the haplotype frequencies (Fig 3b), using the divergence threshold shown in Fig. 3a (estimating 7 haplotypes). Based on the data the true number of haplotypes present in this infection may be as high as 8, suggesting we have captured nearly all of the haplotype diversity at this error tolerance. While in the current analysis, we have been conservative in our treatment of *de novo* mutations and sequencing and/or amplification errors, improvements in laboratory and bioinformatics tools may allow us to distinguish between these categories in the future.

### Relatedness of individual parasites

In addition to providing estimates of the number of distinct haplotypes in an infection, single-cell sequencing can provide details on the patterns of diversity and relatedness contained within each haplotype. We used two common approaches to estimating relatedness between individuals to illustrate this: pairwise allele sharing and identity by descent (IBD). From molecular data, the relatedness of individual parasites can be understood through analysis of sequence identity as well as by contiguous segments of DNA shared between parasites. Long tracts of DNA that contain high identity between any two clones show IBD and have be used to infer the relatedness of individuals in human populations [36, 37]. The fewer the meioses separating two haplotypes, the longer these blocks of IBD will be. Thus, more closely related parasites will share more, and longer, blocks of IBD than unrelated parasites. This process should shape the proportion of alleles two individuals share, though the estimation of allele sharing is not complicated by obstacles such as missing data and variable recombination rates.

Fig. 4a illustrates the proportion of pairwise differences with a UPGMA tree, where highly related individuals cluster together. Based on the threshold suggested above (0.5% SNP differences) 7 unique haplotypes were detected. IBD analysis is broadly concordant with pairwise allele sharing, showing 7 distinct clusters. The mean length and total length of IBD within an infection track closely with pairwise allele sharing, with comparisons between haplotypes with greater numbers of SNP differences also showing smaller blocks of IBD with lower levels of genome-wide IBD (Fig. 4b). The genetic architecture of MAW0 appears to be very similar to what has been seen in polyclonal infections previously collected in Malawi and Thailand[3]. In the future, a larger survey of complex infections may reveal whether this population structure, which includes recent recombinants and more distant lineages, is common or not, and whether other “classes” of complexity exist.

**Fig. 4.**
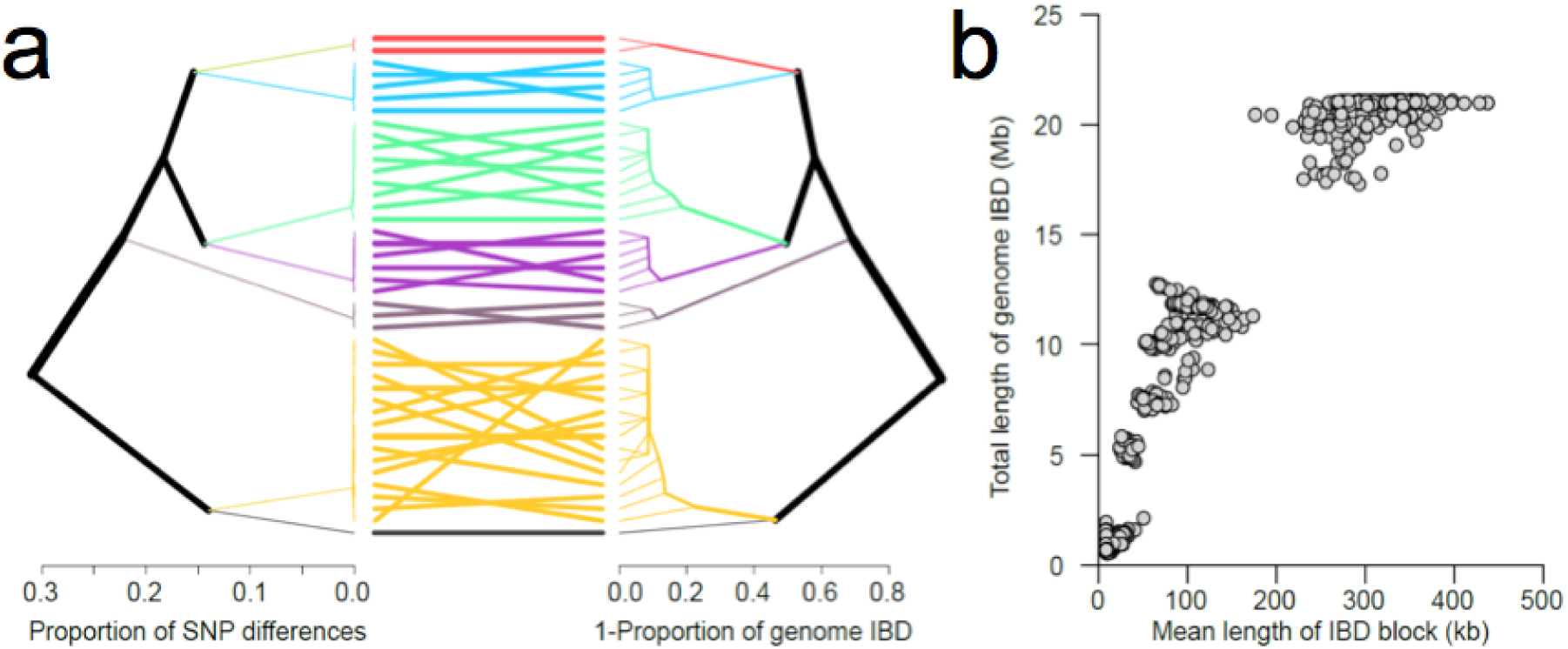
Relatedness of individual parasites. (a) UPGMA tree of pairwise allele sharing (left) and proportion of genome IBD between individual parasites (right) in the MAW0 infection. The haplotypes inferred in Fig 3a are shown in matching colors in the lines joining the tree branches. (b) Relationship between total fraction of IBD and IBD length between parasites. Parasites from identical haplotype groups shared IBD across nearly the entire genome (dots in the upper right), conversely parasites from the most distantly separated haplotype groups (i.e. red vs. dark grey) shared near zero IBD (dots in the bottom left).

### Single-cell genomics accurately captures allele and haplotype frequency in polyclonal infections

This new genome capture strategy includes both extending the time of culture and targeting cells with high DNA content by flow cytometry. Since these actions could place artificial restrictions on which haplotypes are surveyed, it is important to determine whether the single-cell genomes recovered by this method are representative of the diversity found in the original infection. To address this, bulk DNA was extracted from a frozen red blood cell preparation of MAW0. We then compared the allele frequency of 9,766 sites, drawing a comparison between the bulk sample and pooled DNA of 43 out of 48 single-cell genomes passing quality control filtering (Fig. 5). There is high correlation between the datasets (*r*^^2^^=0.96) suggesting minimal sampling bias.

**Fig. 5.**
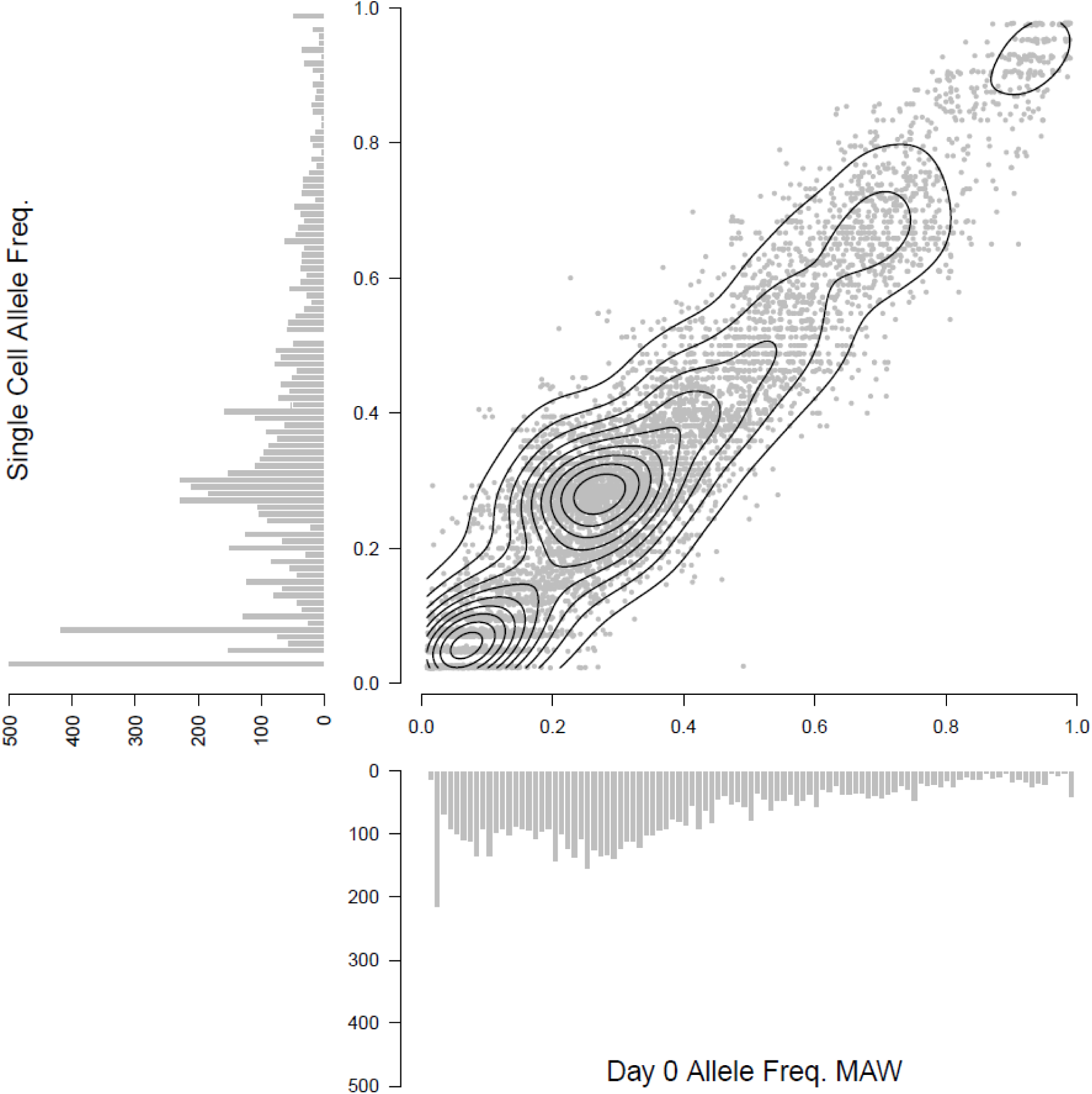
Frequency of alleles detected in bulk DNA at time of thaw and pooled single-cell library data. 9,766 unfixed sites with a read depth of at least 50X in the bulk sample, and had genotype calls for 80% of the single cell sequences were used to estimate sampling bias. A histogram showing the raw counts for each group is attached to the relevant axis. A contour map is overlaid the scatterplot to highlight the density of points lying along the diagonal.

Another way to estimate the sampling bias of single-cell sequencing is to estimate the frequency of each *haplotype* in the bulk sequence data. We can easily determine the prevalence of haplotypes by identifying mutations that are unique to each of the haplotypes. In total 4,375 SNPs were unique to a single haplotype, with a mean of 625 unique SNPs per haplotype. The frequency of each unique SNP and the haplotype it is derived from is shown in Fig. 6. Given this data it was also feasible to correct the abundance of each haplotype in the patient. One haplotype lacked any private mutations, as such its abundance was estimated as the remaining unexplained haplotype frequency (the other inferred haplotype frequencies sum to 0.804). This resulted in a modest improvement in correlation between bulk and single cell allele frequencies to *r*^^2^^=0.98).

**Fig. 6.**
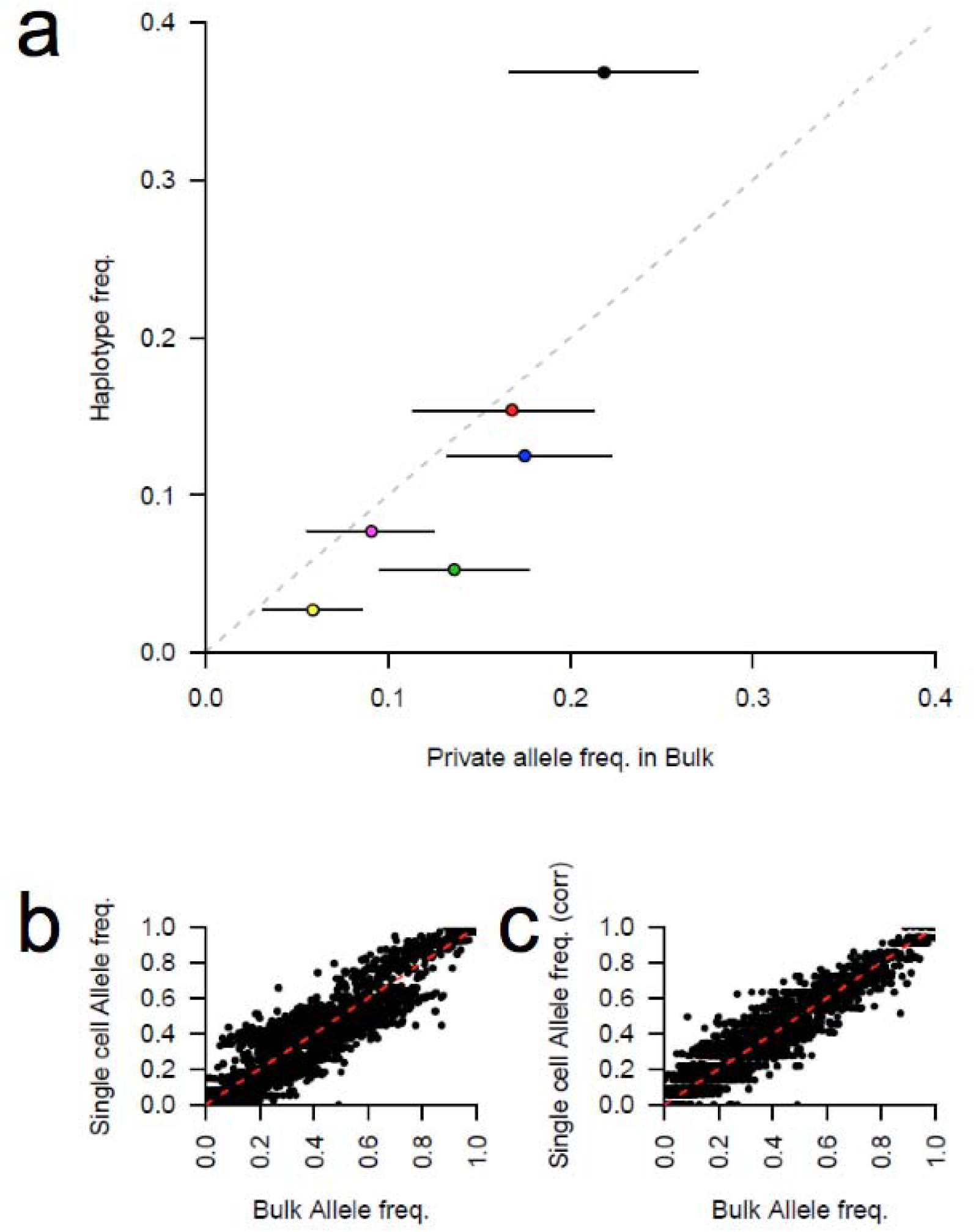
Unique mutations from single-cell sequencing can be used to inferhaplotype abundance in bulk genome sequence. (a) Unique mutations from each haplotype group were used to estimate the bias in estimating their abundance in the single cell sampling. The interquartile range of the allele frequencies for each haplotype is shown by black bars surrounding each plot. These unique allele frequencies were used to correct the haplotype abundances. The original comparison between bulk and single cell allele frequencies ((b); a replicate of Fig. 5) and (c) the corrected data.

A major concern for single-cell genomics is the accurate capture of haplotype diversity and frequency in the original sample. For the complex infection analyzed here, these metrics were maintained. Though this sample suggests 40 hours of culture does not introduce substantial bias, we recommend inclusion of bulk DNA captured at time point zero (directly from the patient arm) for all single-cell genomics analyses as a critical control.

That late-stage parasites, cultured prior to re-invasion may closely capture the abundance of haplotypes found in the original infection is encouraging for future studies. We anticipate this method may be adaptable to the single-cell genome analysis of *Plasmodium vivax*, which cannot currently be cultured for multiple division cycles. Additionally, low-parasitemia infections, which have very small fractions of iRBCs, would be poor samples for flow cytometry due to the likelihood of high false positive rates caused by extended sort times (see Text S1). However, it is possible that magnetic enrichment of late-stage parasites could be performed prior to sorting such that these unknown malaria haplotypes may be individually studied.

## Conclusion

In summary, targeted isolation of late-stage malaria parasites allows efficient, detailed reconstruction of polyclonal malaria infections at the single-cell level. This strategy removed the previous requirement of quality control post-WGA, allowing rapid and cost effective single-cell sequencing to be accessible to most laboratories. Future single-cell genomics approaches may benefit from the strategy of targeting multinucleated cells, especially for Apicomplexan parasites.

## Methods

### Field Sample Collection and Processing

Clinical samples used in this study were obtained from patients presenting to clinics run by the Shoklo Malaria Research Unit in Mae Sot, Thailand, and from a field survey in Chikhwawa, Malawi.

In Malawi, a venous blood sample (5 ml) was collected prior to drug administration from a child aged 47 months presenting to our study site in Chikhwawa with uncomplicated *P. falciparum* malaria (thin smear parasitaemia of 1.4% in 2016). The sample was obtained with the parent’s consent as part of a larger study aimed at understanding within-host parasite genetic diversity in malaria patients from an area of intense malaria transmission. The blood sample was collected in an Acid Citrate Dextrose tube (BD, UK), and transported to the laboratory in Blantyre where it was processed as follows: half of the sample was washed with incomplete RPMI 1640 media and the resulting pellet was mixed with glycerolyte before storage in liquid nitrogen. Parasites used in our FACS experiments were grown from this sample. The other half of the sample was passed through a CF11 column to deplete white blood cells [38] and was stored at -80^^0^^C until needed. Ethical approval for the study was granted by the University Of Malawi, College Of Medicine and Ethics Committee (Protocol number P.02/13/1528) and the Liverpool School of Tropical Medicine Research Ethics Committee (Protocol number 14.035). The *P. falciparum* laboratory line, HB3, used for optimization of gating and WGA experiments was obtained from MR4 (Manassas, VA), and was maintained in the laboratory for several weeks as needed.

### Sterility guidelines

Cell sorting materials and MDA reagents were prepared in “PCR Hood 1” which is housed in a “malaria-DNA free” room separate from the main lab while thawing of frozen single cells and initiation of the MDA protocol was carried out in PCR Hood 2, behind a floor-to-ceiling plastic barrier. MDA was initiated in the main lab on a dedicated thermocycler, while library preparation was performed on a separate thermocycler in another lab. PCR Hood 1 was equipped with standard pipettes and pre-sterile filter tips whereas PCR Hood 2 was equipped with displacement pipettes and pre-sterile displacement tips to reduce the possibility of aerosol contamination between samples. All tubes and tube racks were autoclaved (dry vacuum cycle, 30 min) before use. HB3, THB1 and THB2 cells were sorted into PBS (Qiagen), while MAW0 cells were sorted into recently autoclaved PBS (Lonza).

Prior to use, the interior of the hood was cleaned by wiping down all pipettes, tube racks, and tabletop centrifuges with a series of solutions: 1% bleach, DNAzap according to manufacturer’s instructions, a 70% ethanol wash, and an optional sterile water wash, followed by 15 minutes of UV irradiation. The thermocycler and PCR tube cold rack were wiped down with DNAzap and ethanol before use. We elected not to use UV treatment for reagents and reagent tubes, as the recommended exposure [32] yellowed the manufacturer’s PCR tubes. This may introduce unknown byproducts into the reaction or physically stress the tube, which could compromise sterility.

### Sort preparation

In PCR Hood 1, 5 μl of 1X PBS (AccuGENE) was autoclaved in 400 μl aliquots (in screw cap tubes) or PBS (Qiagen) was delivered into individual, autoclaved PCR tubes with a repeat pipettor. PBS aliquots were stored on a 96 well plate inside of an autoclaved sleeve until use the following day.

### Cell staining

HB3 cells were grown to asynchrony or purified red blood cells isolated from patient samples (∼0.2-0.5 mL) were revived and grown (see Text S2) in resealable culture chambers, flushed with gas (5% CO2, 5% O2, Balance N2). After culture, cells were washed once with PBS and centrifuged (425 x g). 5-8 μl of RBC pellet (∼10^8^ cells, typically) were resuspended in a 1X PBS solution that included 2.5 μl of Vibrant Dye Cycle Green dye(5 mL). The suspension was covered in foil to prevent light exposure and incubated at 37°C for 30 minutes with intermittent inversion every 5-10 minutes. RBCs were washed twice in 10 mL PBS and resuspended in 5-8 mL of PBS and protected from light.

### Cell sorting

Individual cells were sorted into 0.2 mL PCR tubes (5 μl 1X PBS)held on a 96- tube rack one at a time by MoFlo Astrios (Beckman Coulter). Events were gated according to DNA fluorescence and sorted in single-cell sort mode with a drop envelope of 0.5, with each cell typically taking under 15 seconds to sort. Captured cells were immediately stored on dry ice and transferred to -80°C storage.

### Standard PCR

PCR reactions (Takara) included 10 ng of target DNA and 1 μM custom primers (*pfcrt*-L AGGTTCTTGTCTTGGTAAAT and *pfcrt*-R TTTGAATTTCCCTTTTTATT; *dhfr*-F ACGTTTTCGATATTTATGC and *dhfr*-R TCACATTCATATGTACTATTTATTC) using the following program: Hold 940C 2 min; 5 cycles 94^0^C 0.5 min, 50^0^C 0.5 min, 60^0^C 0.5min; 25 cycles 94^0^C 0.5 min, 45^0^C 0.5 min, 60^0^C 0.5 min; Hold 60^0^C 2 min; Hold 4^0^C, and resolved by standard 1% agarose electrophoresis.

### Whole Genome Amplification

Repli-g MDA reagents were thawed, prepared and aliquoted (Qiagen MIDI kit, QIAseq FX single-cell DNA kit) in PCR Hood 1 and transferred to PCR Hood 2. The enzyme mastermix was kept on ice during cell lysing steps. Sorted samples were thawed in PCR Hood 2, spun briefly and the reaction was initiated according to manufacturer’s instructions with the exception of 20 μl of total mastermix added per sample instead of 40 μl for REPLI-g reactions (Text S1). During lysis, tubes were kept outside of the hood on a pre-cooled rack. The reaction proceeded on a thermocycler with heated lid for 4.5 or more hours. We recommend a routine amplification time of 6.5 hours. For 24 MAW0 single-cells, the manufacturer’s protocol for QIAseq FX Single-Cell DNA Kit was followed. In both cases, MDA DNA products were recovered from reaction mixtures with Zymo Genomic Cleanup kits and eluted in 55 μl of water. Library preparations (see below) and MDA products are stored long term at -80C on separate shelves to reduce the potential for contamination.

### Library Preparation KAPA

Illumina sequencing libraries were prepared by the KAPA HyperPlus Kit according to manufacturer’s guidelines using a thermocycler with programmable lid temperature and Agencourt AMPure XP for cleanup and size selection with the following parameters. We used 100 ng of starting DNA material (MDA or bulk extracted DNA), carried out a 1 hr ligation of adapters (5 μl of 15 μm Bioo (NextFlex 48 barcord adapters) in 110 μl reaction), and amplified adapter-ligated libraries for 6 cycles.

### Qiagen

Library preparation was carried out according to manufacturer’s instructions for QIAseq FX single-cell DNA Kit. The recommended standard protocol yielded large, undesired products, so we carried out an additional 1:1 cleanup step followed by an additional size selection step according to the KAPA Hyperplus Kit protocol. Additional experiments revealed improved product distribution by increasing fragmentation incubation time to 33 minutes before proceeding with the recommended cleanup and size-selection step in the Qiagen protocol.

### WGS Library quality control

The size of each Illumina DNA library was determined by HS DNA chips (Agilent) or DNA Tapestation according to manufacturer’s instruction. Pooled libraries were generated by multiplexing either 12 or 24 uniquely barcoded libraries and sequenced on an Illumina HiSeq 2500 using 101bp paired end sequencing with v3 chemistry. Raw sequence reads were de-multiplexed and. fastq files generated using bsl2fastq v2.17.

### Bioinformatics

We aligned each. fastq file to version 3 of the 3D7 reference genome sequence (http://www.plasmodb.org) with BWA-MEM v0.7.5a [39]. PCR duplicates and reads mapping off chromosomal ends were removed with Picard v1.56 (http://broadinstitute.github.io/picard/). We performed base recalibration and realigned around indels using GATK v3.5 [40]. Genotypes were called using GenotypeGVCFs in GATK v3.5 using the QualByDepth, FisherStrand, StrandOddsRatio, VariantType, GCContent and TandemRepeatAnnotator annotations with max_alternate_alleles set to 6. After variant score recalibration we kept all loci with a VQSLOD score > 0 and filtered out SNP calls outside of the “core” genome, defined in [41] For comparative analysis we downsampled bam files to 30X coverage using the - dfrac flag in the GATK engine and calculated coverage statistics using the flagstats and DepthOfCoverage tools. Genomic intervals were subset using Bedtools v2.25.0 [42]

For identity by descent (IBD) analysis we scored regions of IBD using Beagle v4.1 [43]. As this tool was designed for diploid data we generated doubled homozygotes and collapsing together overlapping estimates of IBD. We generated a novel genetic map for this analysis using a collection of genome sequences from clonal Malawian isolates (Nkhoma et al unpublished) using the rhomap function in LDHat v2.2 [44].

All statistical analysis was performed in R v3.3.0 and used the Intervals v0.15.1 [45], R package version 0.15.1 https://CRAN.R-project.org/package=intervals) and SeqArray v1.12.9 [46] SeqArray: Big Data Management of Whole-genome Sequence Variant Calls. R package version 1.12.9. http://github.com/zhengxwen/SeqArray) packages.

## Acknowledgements

Funded by R01 NIH/NIAID AI110941-01A1 to IHC, Wellcome Trust Grant 09999/Z/12/Z to SCN, IHC is a Milton S. and Geraldine M. Goldstein Young Scientist. The Shoklo Malaria Research Unit is part of the Mahidol Oxford University Research Unit, which was supported by the Wellcome Trust of Great Britain. FACS data was generated in the Flow Cytometry Shared Resource Facility which is supported by UTHSCSA, NIH-NCI P30 CA54174 (CTRC at UTHSCSA), and UL1RR025767 (CTSA grant). We thank the patients in Thailand and Malawi, Timothy J.C. Anderson for valuable guidance and Roy Garcia for assistance with sample sequencing.

### Accession Numbers

Raw sequence reads have been deposited to the NCBI short read archive in project PRJNA227205 and [forthcoming].

## Supplementary Material

## Supplementary Tables

Supplementary Table 1. Coverage statistics for all single-cell sequencing experiments. [forthcoming]

## Supplementary Figure Legends

**Fig. S1.**
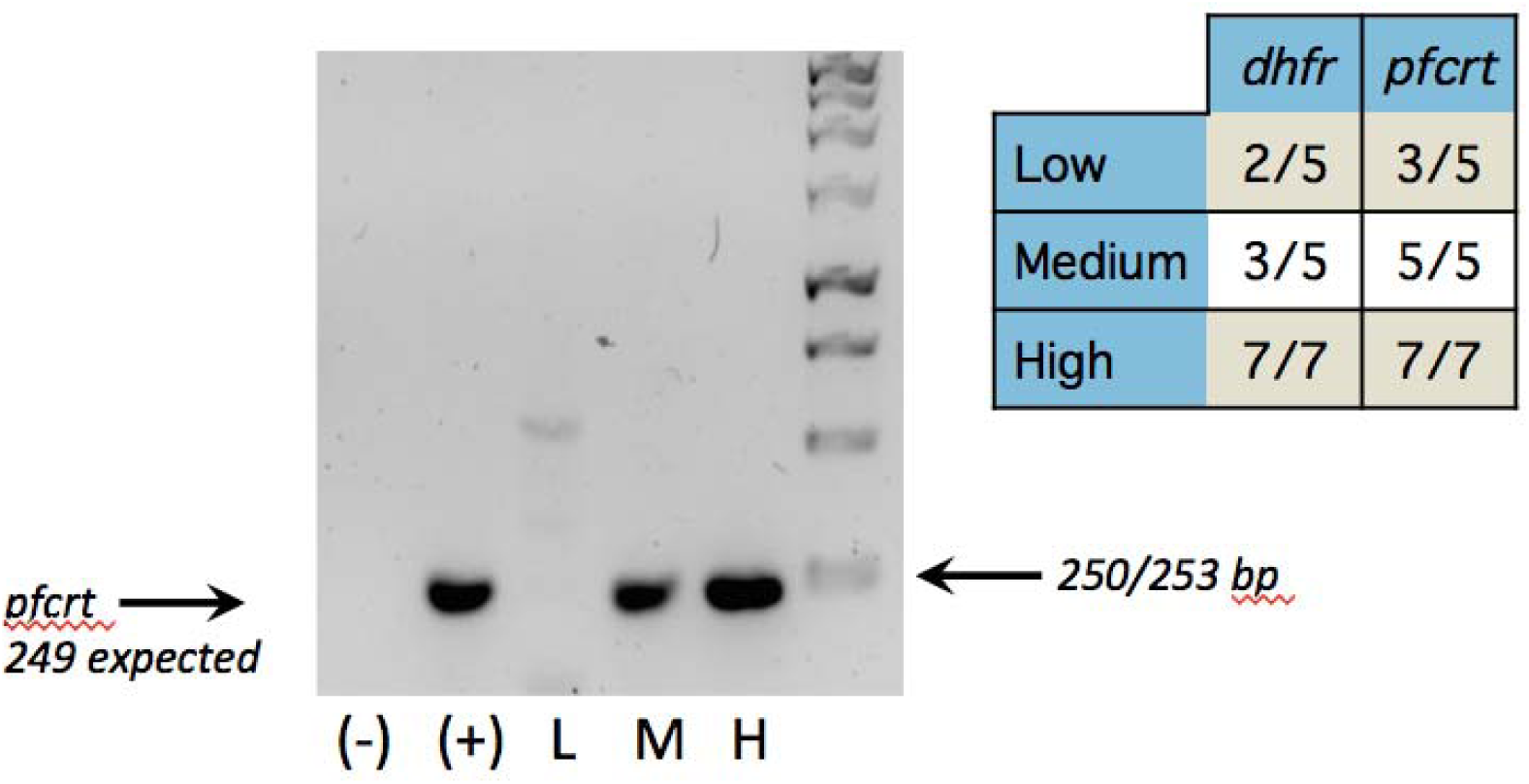
Success rate of *dhfr* and *pfcrt* amplification from L, M, and H gate MDA products. Representative gel for pfcrt end-point PCR product negative control with no DNA (-), HB3 bulk DNA 10 ng (+), MDA product (10 ng) from single-cells captured in L, M, or H gates (L, M, H, respectively) (left). Summary table of successful end-point PCR reactions for each gate and PCR gene product (right).

**Fig. S2.**
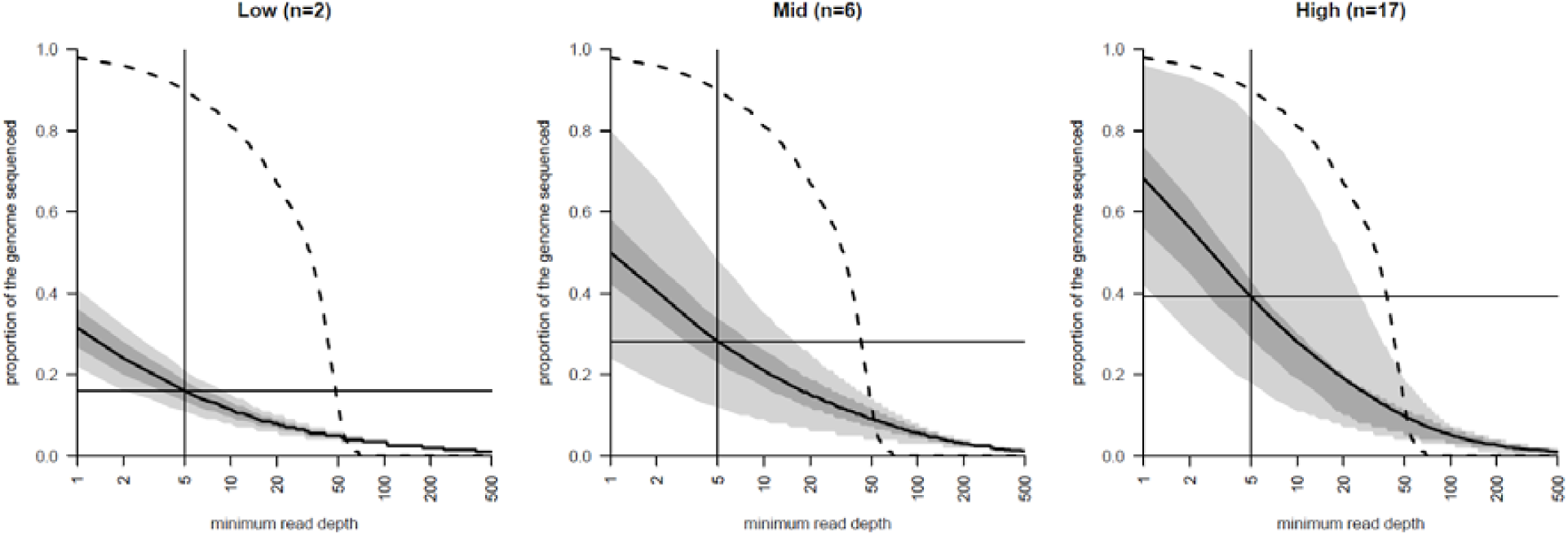
Genome coverage for single-cells collected from the L, M, H gate in two clinical samples (THB1 and THB2). THB1 “M+H” reactions are plotted as “Mid-gate”, as H gate events were infrequent in this sample (see Text S1).

**Fig. S3.**
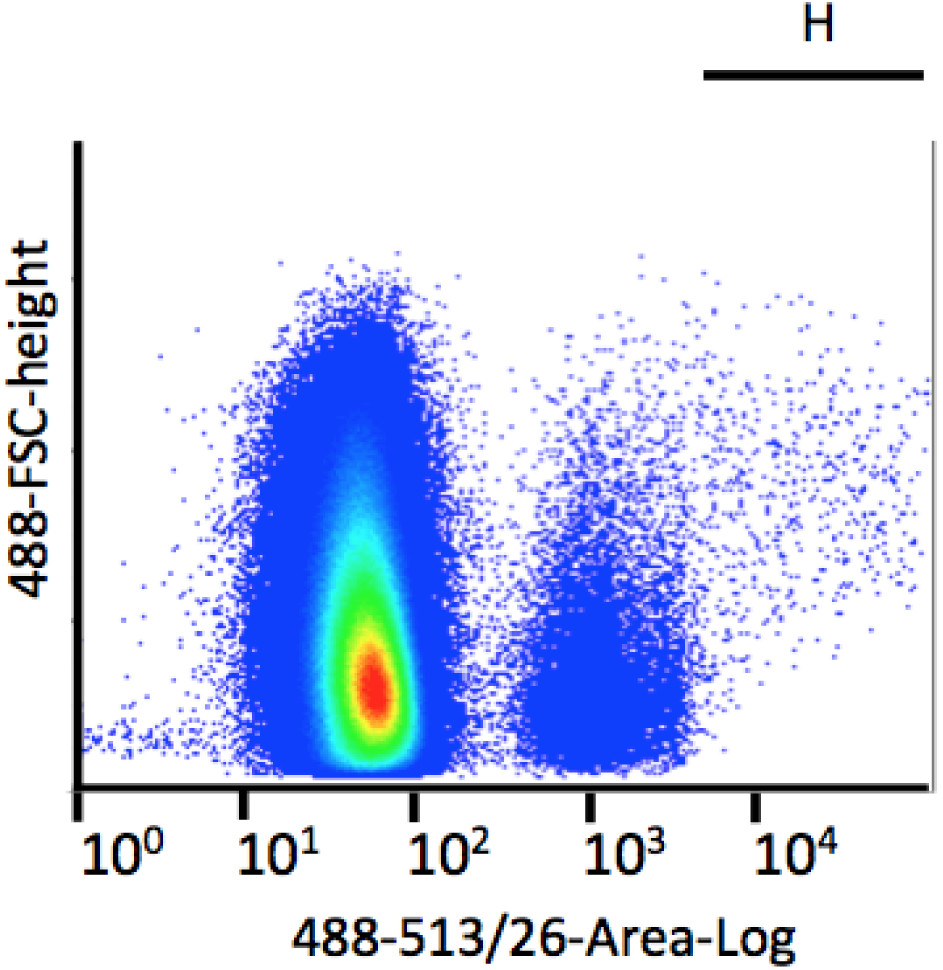
Flow cytometry plot of MAW0. 48 positive events in the H gate were sorted for the single-cell genomics workflow, omitting post-MDA quality control measures.

**Fig. S4.**
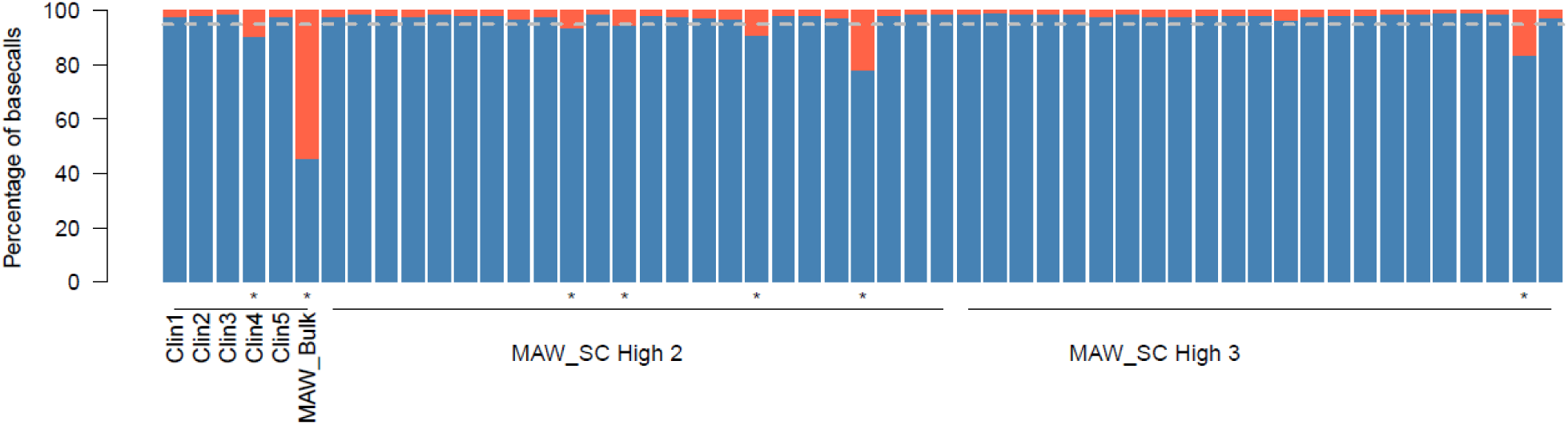
Low rate of contamination in single-cell sequencing data. The proportion of sites with < 95% of reads showing a single genotype call. Sites were filtered to only those likely to be informative (though with a read depth of > 30X). We excluded samples with > 5% unfixed sites, retaining 43 sequences for downstream analysis. Five putatively clonal clinical samples are shown for comparison on the left side of the plot, and the bulk DNA sequence from MAW0.

## Supplementary Text

### Text S1

#### Repli-g MDA optimization

Lowering the reaction time of MDA has been shown to improve genome coverage, likely by restricting the time available for runaway amplification of any given loci to occur. We hypothesized that similar results might be achieved for malaria DNA and sampled single-cell MDA reactions at reaction times of 4.5, 8, or 16 hours prior to deep sequencing. Unexpectedly, we observed similar genome coverage between all reaction times, suggesting these effects may occur reactions times lower than tested. Additionally, we speculate that bias may be reduced in the *P. falciparum* genome due to the prevalence of AT basepairs. One possible mechanism preventing runaway amplification may involve the debranching and re-priming of low-complexity, low-melting point sequences.

In all cases, single-cell MDA REPLI-g reactions generate moderate yields of amplified DNA product (typically ∼0.5-1 μg). With medium-throughput application in mind, we saw a potential opportunity to cut reagent costs by lowering the amount of reaction cocktail used in each sample. In a small comparison of individually-sorted H gate cells (n=2 per sample dilution), similar genome coverage was seen whether using 1X or 0.5 -1.5 μg total yield), but not 0.25X (not detected, <25 ng) of the manufacturer’s recommended reaction cocktail. Thus, subsequent work was carried out using half of the recommended mastermix reaction buffer.

Moving forward, we recommend preparing libraries using QIAseq FX Single Cell Library Kit, which includes MDA and library preparation together, as genomic data quality was higher for samples processed this way.

#### Clinical sample sort optimization

After the initial HB3 experiments, we prepared a clinical sample collected on the Thai-Burmese border using the original protocol to observe whether similar results could be captured from the L, M, and H gates. However, after only 18 hours in culture, a low density of positive events outside of the L gate was observed. This is expected, since the culture time is not long enough to allow for progression to later stages of the cell cycle. Thus, only two flow cytometry gates were collected: L as well as the M and H gates combined. We additionally did not pre-screen samples for quality prior to library amplification. In this experiment, 7 of 10 tested cells in the M and H combined gate had 6% or less of reads map to the reference genome (Table S1), while 7 of 10 cells from the L gate had > =80% reads map. Thus, L gate sorted cells had a much lower chance of environmental contamination than the M and H combined gate. Since early-stage cells dominated the culture at that time point, events in the M and H gates were infrequent and took longer to sort. We hypothesize that the increased time for which the tube was exposed to ambient air increased the likelihood of contamination. Alternatively, observed events in the M and H combined gates could have included low-frequency false positive machine artifacts. Additionally, the mean genome coverage of the 3 successful library preparations sorted by the M and H combined gate was higher than the genome coverage observed by 7 L gate-sorted cells, consistent with the trend observed in HB3.

We reasoned it may be possible to increase the frequency of events in the M and H gates by growing the samples for 40 hours, instead of for 18 hours. Indeed, for a second sample collected in the same region (THB2) and grown for this extended period, events were more dense in the M and H gate, which brought down the time to sort individual cells. For this and other experiments, successful events were generally sorted in < 15 s per sample, though we note the window for avoiding contamination may vary substantially from lab to lab. In all cases for THB2, read purity was > 94% in all gates. Furthermore, we observed a trend in improved genome coverage similar to HB3 data, where H gate cells generated roughly twice as much coverage as L or M gate cells (Table S1).

Finally, the genome coverage observed for cells in the THB2 H gate is less than what is observed for MAW0. Between these two experiments, we switched from using the manufacturer’s PBS to in-house autoclaved AccuGENE 1X PBS (Lonza) for the sort capture buffer (5 μl in a single 0.2 mL PCR tube). These and additional experiments suggest that using recently autoclaved AccuGENE 1X PBS (Lonza) in place of the PBS provided by Qiagen may contribute to increased quality metrics.

### Text S2

#### PROTOCOL

#### REAGENTS & SOLUTIONS

Vibrant Dye Cycle Green (#V35004)

Incomplete cell media (ICM)- 500 mL RPMI 1640 (Gibco #11875119), 12.5 mL

HEPES (Gibco #15630-080), 1 mL Gentamicin 10 mg/mL (Gibco #15719-064)

Complete cell media (CM), ICM 313 mL, 25 g AlbumaxII (Thermo #11021-029), 0.156g Hypoxanthine (Sigma #H936)

10X PBS (Ambion #AM9624)

NaCl solutions, NaCl (Sigma #S-7653) in sterile water (Gibco #15230162), filtered by 500 mL filter system (Corning #430770)

Culture flasks (Corning #430168)

AccuGENE 1X PBS (Lonza #51225)

DNAZap (Thermo Scientific #AM9890)

Bleach (Essendant #KIKBLEACH6)

Sterile water (Gibco #15230196)

Nalgene PETG erlenmeyer flasks (Thermo Scientific #41120250)

Free-Standing Microcentrifuge Tubes with Screw Caps (Fisherbrand #02-682- 558)

REPLI-g Midi kit (Qiagen #150045)

KAPA Hyperplus Library Kit with library amplification (KAPA #KK8514)

Bioo NEXTflex DNA Barcodes (#514104)

QIAseq FX single-cell RNA library kit (Qiagen #180733) Agencourt AMPure XP (Beckman #A63882)

80% ethanol (Fisher Bioreagents)

Glycerolyte (Fenwal #4A7831)

NEXTflex 48 barcodes (Bioo #514104)

Genomic DNA Clean & Concentrator-10 (Zymo #D4010)

#### EQUIPMENT

PCR Workstation (Airclean 600) “PCR HOOD #1” & “PCR HOOD #2”

Tabletop LSE microcentrifuge (Corning #6765)

Thermocycler 1 (GeneAmp PCR System 9700 Thermo)

Thermocycler 2 (PTC-200 MJ Reseearch)

Incubator Culture Chamber (C.B.S. # M-624)

PCR cold-rack (Eppendorf #022510509)

Microman M10 positive displacement pipette (#F148501G)

Microman M100 positive displacement pipette (#F148504G)

Gilson CP10ST Microman tips (#F148413G)

Gilson CP100ST Microman tips (#F148415G)

Finnpipette Novus Electronic Single-Channel Pipette (Thermo Scientific #9400250)

PCR tubes (Phenix Research #MPX-200)

Sterilization Pouches (Fisherbrand #01-812-54)

Magnetic Stand (Ambion #AM10027)

Cardinal Health Secure-Gard Cone Mask (Dupont Personal Protection #AT7509)

Sterile gowns (Kimberly-Clark #90042)

#### FILTER HOOD CLEANING PROCEDURE

Don sterile gowns, new gloves, and cone mask.

Prepare 1% bleach in sterile water.

Using a dropper, wet sterile paper towel (included in sterile gown package) with 1% bleach and wipe internal surfaces of PCR HOOD. Dry surfaces with a clean paper towel.

Use DNAzap according to manufacturer’s instructions, including all surfaces and pipettes.

Spray and wipe surfaces with 70% ethanol.

Wipe surfaces with sterile water.

Turn UV light on for 15 minutes.

Unwrap sterile tips and PETG flask (for waste) in hood without touching any surfaces.

Wipe the outside surface of all reagents with 70% ethanol prior to use in the hood.

#### CELL CULTURE

Thaw purified RBCs at 37^^0^^C for 1-2 minutes.

Add 12% NaCl (1/5th volume of RBC) dropwise while swirling sample. Let stand for 5 minutes.

Add 1.8% NaCl (5 mL) dropwise while swirling sample. Let stand for 2 min.

Add 0.9% NaCl (5 mL) dropwise while swirling sample. Let stand for 2 min.

Wash cells once in 10 mL ICM, using centrifugation at 425 x g for 5 min.

Add 8 mL CM grow in sealed box flushed with 5% CO2 5% O2, balance N2 at 37C for 40 hr.

#### SORT TUBE PREPARATION

In PCR HOOD #1, seal approximately two hundred 0.2 mL PCR tubes, two PCR tube plates and several aliquots of 1X PBS (Lonza) in sterilization pouches.

Autoclave on dry vacuum program (30 minutes).

Clean PCR HOOD#1.

Unwrap sterile PETG flask and pre-sterile 200 μl filter tips.

Wipe outside of post-autoclave sterilization pouches with 70% ethanol.

Dispense 5 μl of autoclaved PBS into individual 0.2 mL PCR tubes using a repeat pipettor.

Store tubes at RT overnight on racks, with each rack protected in its sterilization pouch (opened but folded over to prevent air flow).

#### STAINING

Wash culture 1X times in PBS by centrifugation (425 x g)

Freeze aliquot of resultant pellet for bulk DNA library preparation (typically 100 μl of pellet).

Add 7-8 μL of pellet to 5 mL staining buffer (1X PBS, 2.5 μL Vibrant DyeCycle Green). Protect tube from light with foil.

Incubate at 37°C with intermittent inversion every 5 minutes for 30 minutes.

Wash cells twice in 1X PBS.

Resuspend pellet in 5 mL 1X PBS.

MDA + LIBRARY PREPARATION (*preferred method*) Qiagen

Clean PCR HOOD #2. Cool PCR cold-rack on ice next to PCR HOOD #2.

Clean thermocycler with DNAzap and 70% ethanol. Stock PETG flask for waste.

Thaw MDA reagents in PCR HOOD #2. We routinely processed 24 reactions at once.

Place MasterMix on ice near PCR HOOD #2, place D2 and Stop solutions in

PCR HOOD #2.

Thaw captured cells at RT, pulse on tabletop centrifuge inside of PCR Hood #2.

Use the PCR cold-rack outside of PCR HOOD #2 for “on ice” incubations, to minimize contact of the tube with ice.

Follow the QIAseq FX single-cell DNA Kit manufacturer’s instructions for MDA.

Elute DNA with 55 μl dH2O.

Quantify DNA with Qubit BR Assay kit (#Q32850), according to manufacturer’s instructions

#### Clean-up

Purify MDA DNA products with Genomic DNA Clean & Concentrator-10

(according to manufacturer’s instructions, using 14,000 x g for centrifugation steps.

#### Library Prep

Follow the QIAseq FX single-cell DNA Kit manufacturer’s instructions for PCRfree library preparation. We included the optional enhancer. Elute DNA with 38 μl of dH2O. Typical concentrations of prepped libraries were 3-8 ng/βl. Additional experiments revealed improved product distribution by increasing fragmentation incubation time to 33 minutes before proceeding with the recommended cleanup and size-selection step in the Qiagen protocol.

### WHOLE GENOME AMPLIFICATION (*intermediate method*) (REPLI-G, Qiagen)

Follow manufacturer’s guidelines with the following additions:

*Preparation*

Autoclave 2 mL tubes (3).

Clean PCR HOOD #1 and #2. Cool PCR cold-rack on ice next to PCR HOOD #2.

Clean thermocycler with DNAzap and 70% ethanol.

Prepare D2, MasterMix, and aliquot Stop stocks in PCR HOOD #1. MasterMix was prepared at half volume. We routinely processed 24 reactions at once.

Place MasterMix on ice near PCR HOOD #2, place D2 and Stop solutions in PCR HOOD #2.

Thaw captured cells at RT, pulse on tabletop centrifuge inside of PCR Hood #2.

Follow Qiagen REPLI-g protocol, using the PCR cold-rack outside of PCR HOOD #2 for “on ice” incubations, to minimize contact of the tube with ice.

Deliver half of the recommended volume of MasterMix per sample (20 μl).

Incubate at 30°C for 6.5 hours, followed by a 3 min incubation at 65°C for reaction inhibition. Can hold overnight at 4°C.

WGS LIBRARY PREPARATION (*intermediate method, continued*) KAPA

Follow the KAPA Hyperplus Library Amplification Kit manufacturer’s instructions with the following parameters:

Initiate each reaction with 100 ng purified MDA DNA.

Carry fragmentation out for 25 minutes.

Use 5 μl of 15 μM illumina-compatible adapters (Bioo).

Use 6 cycles total for PCR amplification.

Elute DNA with 38 μl of dH2O. Typical concentrations of prepped libraries were 5-10 ng/μl.

## References

1. Wang J, Song Y: Single cell sequencing: a distinct new field. Clin Transl Med 2017, 6:10.

2. Gawad C, Koh W, Quake SR: Single-cell genome sequencing: current state of the science. Nat Rev Genet 2016, 17:175–188.

3. Nair S, Nkhoma SC, Serre D, Zimmerman PA, Gorena K, Daniel BJ, Nosten F, Anderson TJ, Cheeseman IH: Single-cell genomics for dissection of complex malaria infections. Genome Res 2014, 24:1028–1038.

4. Blake DP, Clark EL, Macdonald SE, Thenmozhi V, Kundu K, Garg R, Jatau ID, Ayoade S, Kawahara F, Moftah A, et al: Population, genetic, and antigenic diversity of the apicomplexan Eimeria tenella and their relevance to vaccine development. Proc Natl Acad Sci U S A 2015, 112:E5343–5350.

5. Navin N, Kendall J, Troge J, Andrews P, Rodgers L, McIndoo J, Cook K, Stepansky A, Levy D, Esposito D, et al: Tumour evolution inferred by single-cell sequencing. Nature 2011, 472:90–94.

6. Lodato MA, Woodworth MB, Lee S, Evrony GD, Mehta BK, Karger A, Lee S, Chittenden TW, D’Gama AM, Cai X, et al: Somatic mutation in single human neurons tracks developmental and transcriptional history. Science 2015, 350:94–98.

7. Miotto O, Amato R, Ashley EA, MacInnis B, Almagro-Garcia J, Amaratunga C, Lim P, Mead D, Oyola SO, Dhorda M, et al: Genetic architecture of artemisinin-resistant Plasmodium falciparum. Nat Genet 2015, 47:226–234.

8. Manske M, Miotto O, Campino S, Auburn S, Almagro-Garcia J, Maslen G, O’Brien J, Djimde A, Doumbo O, Zongo I, et al: Analysis of Plasmodium falciparum diversity in natural infections by deep sequencing. Nature 2012, 487:375–379.

9. Amambua-Ngwa A, Tetteh KK, Manske M, Gomez-Escobar N, Stewart LB, Deerhake ME, Cheeseman IH, Newbold CI, Holder AA, Knuepfer E, et al: Population genomic scan for candidate signatures of balancing selection to guide antigen characterization in malaria parasites. PLoS Genet 2012, 8:e1002992.

10. Conway DJ, Greenwood BM, McBride JS: The epidemiology of multiple-clone Plasmodium falciparum infections in Gambian patients. Parasitology 1991, 103 Pt 1:1–6.

11. Conway DJ, McBride JS: Population genetics of Plasmodium falciparum within a malaria hyperendemic area. Parasitology 1991, 103 Pt 1:7–16.

12. Nkhoma SC, Nair S, Cheeseman IH, Rohr-Allegrini C, Singlam S, Nosten F, Anderson TJ: Close kinship within multiple-genotype malaria parasite infections. Proc Biol Sci 2012, 279:2589–2598.

13. Snounou G, Viriyakosol S, Zhu XP, Jarra W, Pinheiro L, do Rosario VE, Thaithong S, Brown KN: High sensitivity of detection of human malaria parasites by the use of nested polymerase chain reaction. Mol Biochem Parasitol 1993, 61:315–320.

14. Nkhoma SC, Nair S, Al-Saai S, Ashley E, McGready R, Phyo AP, Nosten F, Anderson TJ: Population genetic correlates of declining transmission in a human pathogen. Mol Ecol 2013, 22:273–285.

15. Hill WG, Babiker HA: Estimation of numbers of malaria clones in blood samples. Proc Biol Sci 1995, 262:249–257.

16. Galinsky K, Valim C, Salmier A, de Thoisy B, Musset L, Legrand E, Faust A, Baniecki ML, Ndiaye D, Daniels RF, et al: COIL: a methodology for evaluating malarial complexity of infection using likelihood from single nucleotide polymorphism data. Malar J 2015, 14:4.

17. Juliano JJ, Porter K, Mwapasa V, Sem R, Rogers WO, Ariey F, Wongsrichanalai C, Read A, Meshnick SR: Exposing malaria in-host diversity and estimating population diversity by capture-recapture using massively parallel pyrosequencing. Proc Natl Acad Sci U S A 2010, 107:20138–20143.

18. O’Brien JD, Iqbal Z, Wendler J, Amenga-Etego L: Inferring Strain Mixture within Clinical Plasmodium falciparum Isolates from Genomic Sequence Data. PLoS Comput Biol 2016, 12:e1004824.

19. Assefa SA, Preston MD, Campino S, Ocholla H, Sutherland CJ, Clark TG: estMOI: estimating multiplicity of infection using parasite deep sequencing data. Bioinformatics 2014, 30:1292–1294.

20. Pearson RD, Amato R, Auburn S, Miotto O, Almagro-Garcia J, Amaratunga C, Suon S, Mao S, Noviyanti R, Trimarsanto H, et al: Genomic analysis of local variation and recent evolution in Plasmodium vivax. Nat Genet 2016, 48:959–964.

21. Zhu S, Almagro-Garcia J, McVean G: Deconvolution of multiple infections in Plasmodium falciparum from high throughput sequencing data. 2017.

22. Song S, Sliwerska E, Emery S, Kidd JM: Modeling Human Population Separation History Using Physically Phased Genomes. Genetics 2017, 205:385–395.

23. Rosario V: Cloning of naturally occurring mixed infections of malaria parasites. Science 1981, 212:1037–1038.

24. Leung ML, Wang Y, Waters J, Navin NE: SNES: single nucleus exome sequencing. Genome Biol 2015, 16:55.

25. Oyola SO, Ariani CV, Hamilton WL, Kekre M, Amenga-Etego LN, Ghansah A, Rutledge GG, Redmond S, Manske M, Jyothi D, et al: Whole genome sequencing of Plasmodium falciparum from dried blood spots using selective whole genome amplification. Malar J 2016, 15:597.

26. Reilly HB, Wang H, Steuter JA, Marx AM, Ferdig MT: Quantitative dissection of clone-specific growth rates in cultured malaria parasites. Int J Parasitol 2007, 37:1599–1607.

27. Dekel E, Rivkin A, Heidenreich M, Nadav Y, Ofir-Birin Y, Porat Z, Regev-Rudzki N: Identification and classification of the malaria parasite blood developmental stages, using imaging flow cytometry. Methods 2017, 112:157–166.

28. Gole J, Gore A, Richards A, Chiu YJ, Fung HL, Bushman D, Chiang HI, Chun J, Lo YH, Zhang K: Massively parallel polymerase cloning and genome sequencing of single cells using nanoliter microwells. Nat Biotechnol 2013, 31:1126–1132.

29. Suyama T, Kawaharasaki M: Decomposition of waste DNA with extended autoclaving under unsaturated steam. Biotechniques 2013, 55:296–299.

30. Motley ST, Picuri JM, Crowder CD, Minich JJ, Hofstadler SA, Eshoo MW: Improved multiple displacement amplification (iMDA) and ultraclean reagents. BMC Genomics 2014, 15:443.

31. Gardner MJ, Hall N, Fung E, White O, Berriman M, Hyman RW, Carlton JM, Pain A, Nelson KE, Bowman S, et al: Genome sequence of the human malaria parasite Plasmodium falciparum. Nature 2002, 419:498–511.

32. Woyke T, Sczyrba A, Lee J, Rinke C, Tighe D, Clingenpeel S, Malmstrom R, Stepanauskas R, Cheng JF: Decontamination of MDA reagents for single cell whole genome amplification. PLoS One 2011, 6:e26161.

33. Claessens A, Hamilton WL, Kekre M, Otto TD, Faizullabhoy A, Rayner JC, Kwiatkowski D: Generation of antigenic diversity in Plasmodium falciparum by structured rearrangement of Var genes during mitosis. PLoS Genet 2014, 10:e1004812.

34. de Bourcy CF, De Vlaminck I, Kanbar JN, Wang J, Gawad C, Quake SR: A quantitative comparison of single-cell whole genome amplification methods. PLoS One 2014, 9:e105585.

35. Colwell RK, Chao A, Gotelli NJ, Lin SY, Mao CX, Chazdon RL, Longino JT: Models and estimators linking individual-based and sample-based rarefaction, extrapolation and comparison of assemblages. Journal of Plant Ecology 2012, 5:3–21.

36. Ralph P, Coop G: The geography of recent genetic ancestry across Europe. PLoS Biol 2013, 11:e1001555.

37. Gusev A, Palamara PF, Aponte G, Zhuang Z, Darvasi A, Gregersen P, Pe’er I: The architecture of long-range haplotypes shared within and across populations. Mol Biol Evol 2012, 29:473–486.

38. Venkatesan M, Amaratunga C, Campino S, Auburn S, Koch O, Lim P, Uk S, Socheat D, Kwiatkowski DP, Fairhurst RM, Plowe CV: Using CF11 cellulose columns to inexpensively and effectively remove human DNA from Plasmodium falciparum-infected whole blood samples. Malar J 2012, 11:41.

39. Li H: Aligning sequence reads, clone sequences and assembly contigs with BWA-MEM. arXivorg 2013.

40. DePristo MA, Banks E, Poplin R, Garimella KV, Maguire JR, Hartl C, Philippakis AA, del Angel G, Rivas MA, Hanna M, et al: A framework for variation discovery and genotyping using next-generation DNA sequencing data. Nat Genet 2011, 43:491–498.

41. Miles A, Iqbal Z, Vauterin P, Pearson R, Campino S, Theron M, Gould K, Mead D, Drury E, O’Brien J, et al: Indels, structural variation, and recombination drive genomic diversity in Plasmodium falciparum. Genome Res 2016, 26:1288–1299.

42. Quinlan AR: BEDTools: The Swiss-Army Tool for Genome Feature Analysis. Curr Protoc Bioinformatics 2014, 47:11-12 11-34.

43. Browning BL, Browning SR: Improving the accuracy and efficiency of identity-by-descent detection in population data. Genetics 2013, 194:459–471.

44. Auton A, McVean G: Recombination rate estimation in the presence of hotspots. Genome Res 2007, 17:1219–1227.

45. Bourgon R: intervals: Tools for Working with Points and Intervals. 2015.

46. Zheng X, Gogarten S: SeqArray: Big Data Management of Whole-genome Sequence Variant Calls. 2016.

